# Deciphering the Biological Function of Tandem Mass Spectra Using a Dual-Modal Metabolite Language Model

**DOI:** 10.64898/2026.11.16.766993

**Authors:** Feifan Zhang, Yuting Wang, Yijiang Liu, Jiramet Kinchagawat, Chuchu Wang, Wenya Wang, Yifan Fu, Qingpeng Kong, Xiaotao Shen

## Abstract

Liquid chromatography-tandem mass spectrometry untargeted metabolomics captures thousands of metabolic peaks from biological samples. Conventional identification-dependent metabolomics workflows require confident structural annotation before identified features can enter downstream pathway or functional analysis. However, only a small fraction of detected metabolic features can be assigned confident structural annotations, leaving a large fraction difficult to incorporate into downstream biological interpretation, a form of ‘functional dark matter’. Here, we introduce Spec2Function, a function-centric framework that infers biological functions from tandem mass spectrometry (MS^2^) spectra. Spec2Function is trained using characterized spectrum-text pairs but does not require prior structural identification of the query spectra at inference time. At its core is Spec2Biotext, a dual-encoder contrastive model that embeds MS^2^ spectra and curated multi-aspect natural language functional texts into a unified functional space. Combined with a retrieval-augmented generation framework grounded in curated databases and biomedical literature, Spec2Function retrieves evidence-grounded biological knowledge and produces coherent functional summaries for individual spectra or groups of spectra. Across metabolite- and pathway-level benchmarks, Spec2Function achieved higher similarity to reference functional descriptions than the evaluated comparators and expanded the fraction of the metabolome accessible to functional interpretation. Application to a human liver cirrhosis cohort recapitulated known metabolic alterations and identified additional literature-supported functional modules. Applying Spec2Function to a 30-organ macaque aging atlas revealed organ-specific metabolic functions and previously unreported functional modules that were supported by independent human and mouse datasets. Together, these results position Spec2Function as a function-centric framework for scalable biological interpretation of structurally unannotated MS^2^ spectra.

## Introduction

Liquid chromatography-tandem mass spectrometry (LC-MS/MS) untargeted metabolomics enables comprehensive profiling of metabolites across biological samples, offering powerful insights into cellular physiology, disease mechanisms, and environmental responses^1–4^. Each detected metabolic peak, represented as a retention time (RT) and mass-to-charge (*m/z*) pair, typically corresponds to a putative biochemical entity^1^. However, the functional interpretation of this vast pool of peaks remains a major challenge^5,6^.

Conventional identification-dependent LC-MS/MS metabolomics workflows typically first assign structural annotations and subsequently perform pathway or functional analysis on the identified subset^7–9^. However, accurate structure identification remains a major bottleneck in this field^5,10^. Most identification approaches rely on matching MS^2^ spectra (generated by fragmenting precursor ions) to reference spectral libraries^11–13^, which are incomplete, unevenly curated, and biased toward well-characterized metabolites^9,11^. Structural isomers, instrument-specific fragmentation variability, and the absence of standardized retention time information further complicate reliable structural identification^5,14^. Consequently, only a small proportion of detected metabolic features can be assigned confident structural annotations, leaving a large fraction of features difficult to incorporate into downstream biological interpretation and hypothesis generation^11^.

Although MS^2^ spectra are traditionally used for structural identification, they also encode rich fragmentation patterns that may carry information associated with underlying biological roles^15–17^. This raises an important question: can MS^2^ spectra be used to infer biological functions for experimentally observed metabolic features before confident structural identification? Several efforts suggest the answer is yes^18–20^. For instance, GNPS leverages MS^2^ spectral similarity networking to cluster peaks into molecular families, enabling unannotated peaks to be associated with characterized ones and thereby facilitating functional inference^20^. Similarly, MS2MP employs deep learning to predict the association between MS^2^ spectra and KEGG pathways, allowing indirect inference of biological roles of peaks^19^. Collectively, these studies demonstrate that MS^2^ spectra can support biological function inference for metabolic features that remain structurally unannotated in untargeted experiments.

However, existing approaches remain fragmented, and a general framework for systematically linking MS^2^ spectra to multi-aspect biological functional information remains lacking. A central obstacle is the inherent modality gap. MS^2^ spectra are encoded as numerical vectors or fragmentation graphs^21–23^, whereas metabolite functions are typically described in natural language^24,25^. This modality gap has historically limited direct mapping between spectral data and biological function.

Recent breakthroughs in large language models (LLMs) have revolutionized biological data modeling^26–29^, opening new avenues for representing and reasoning over complex, multimodal inputs^30,31^. MS^2^ spectra can be transformed into token-based representations that are compatible with deep learning architectures^15,21^. At the same time, curated databases such as HMDB^25^ and KEGG^24^ provide rich textual descriptions of metabolite functions across physiological and disease contexts, serving as valuable sources of functional semantics, which we refer to here as biological text (biotext). Importantly, recent progress in cross-modal contrastive learning has enabled the joint embedding of heterogeneous data types across diverse biomedical applications^32–35^. Extending this idea further, embedding both MS^2^ spectra and biological function texts (biotext) into a shared representation space offers a promising strategy for directly inferring metabolite functions when the corresponding metabolic features remain structurally unannotated in untargeted experiments.

Based on this concept, we developed Spec2Function, a cross-modal framework that infers biological function from MS^2^ spectra. Spec2Function is trained using characterized spectrum-text pairs, but does not require prior structural identification of a query spectrum at inference time. At the core of Spec2Function is a dual-encoder model, Spec2Biotext, which consists of a spectral encoder that transforms MS^2^ spectra into functionally informative vector representations and a biotext encoder that converts biological functional text into structurally aware functional embeddings. The two encoders are co-trained using a contrastive learning objective, driving MS^2^ spectra and their associated biotext descriptions to align within a shared functional space. After training, this dual-modal model (Spec2Biotext) allows MS^2^ spectra to be queried against biotext representations, enabling the retrieval of relevant functional information. Spec2Function supports multiple downstream applications. For a single MS^2^ spectrum, it retrieves the most functionally similar biotexts and summarizes the inferred biological roles using an LLM-based functional inference module with retrieval-augmented generation (RAG)^36^. For differential metabolomics studies, it aggregates functionally similar biotexts for a set of MS^2^ spectra and performs set-level functional analysis to uncover biological functions^28,37^.

Systematic benchmarking against an identification-dependent pathway-analysis workflow and complementary structure-, spectrum-, and pathway-based approaches showed that Spec2Function extends functional analysis to a broader fraction of disease-associated MS^2^ features while recovering biologically coherent modules, including signals not recovered by the identification-dependent analysis. To further evaluate its utility in real-world applications, we applied Spec2Function to a multi-organ macaque (*Macaca mulatta*) aging dataset. This analysis enabled cross-tissue functional profiling and revealed both conserved and organ-specific metabolic signatures of aging, including previously unrecognized functional modules that were subsequently validated across species. Based on these results, we constructed a comprehensive multi-organ metabolic aging atlas of the female macaque (Macaque Multi-Organ Metabolic Aging Atlas, MOMA), which is available at https://moma.jaspershenlab.com/.

Taken together, Spec2Function provides a function-centric framework that enables structurally unannotated metabolic features to be incorporated into functional analysis before confident structural identification is available. This function-centric strategy expands the usable metabolome, providing a scalable and evidence-grounded route for extracting biological insights from the vast and largely unannotated landscape of untargeted metabolomics.

## Results

### Spec2Biotext: a dual-encoder model that aligns MS^2^ spectra with biological function

Traditional LC-MS/MS untargeted metabolomics follows an identification-dependent framework (MS^2^ spectra-based annotations and pathway analysis), limiting downstream biological interpretation (Fig. 1a). This underscores the need for a function-centric framework that infers biological meaning from MS^2^ spectra without requiring prior structural identification of the query spectrum at inference time (Fig. 1a). Achieving this requires a model that aligns two disparate modalities: MS^2^ spectra and biotext describing metabolite function. We therefore developed Spec2Biotext, a multimodal dual-encoder model that learns a shared representation between MS^2^ spectra and biological function descriptions.

**Fig. 1:**
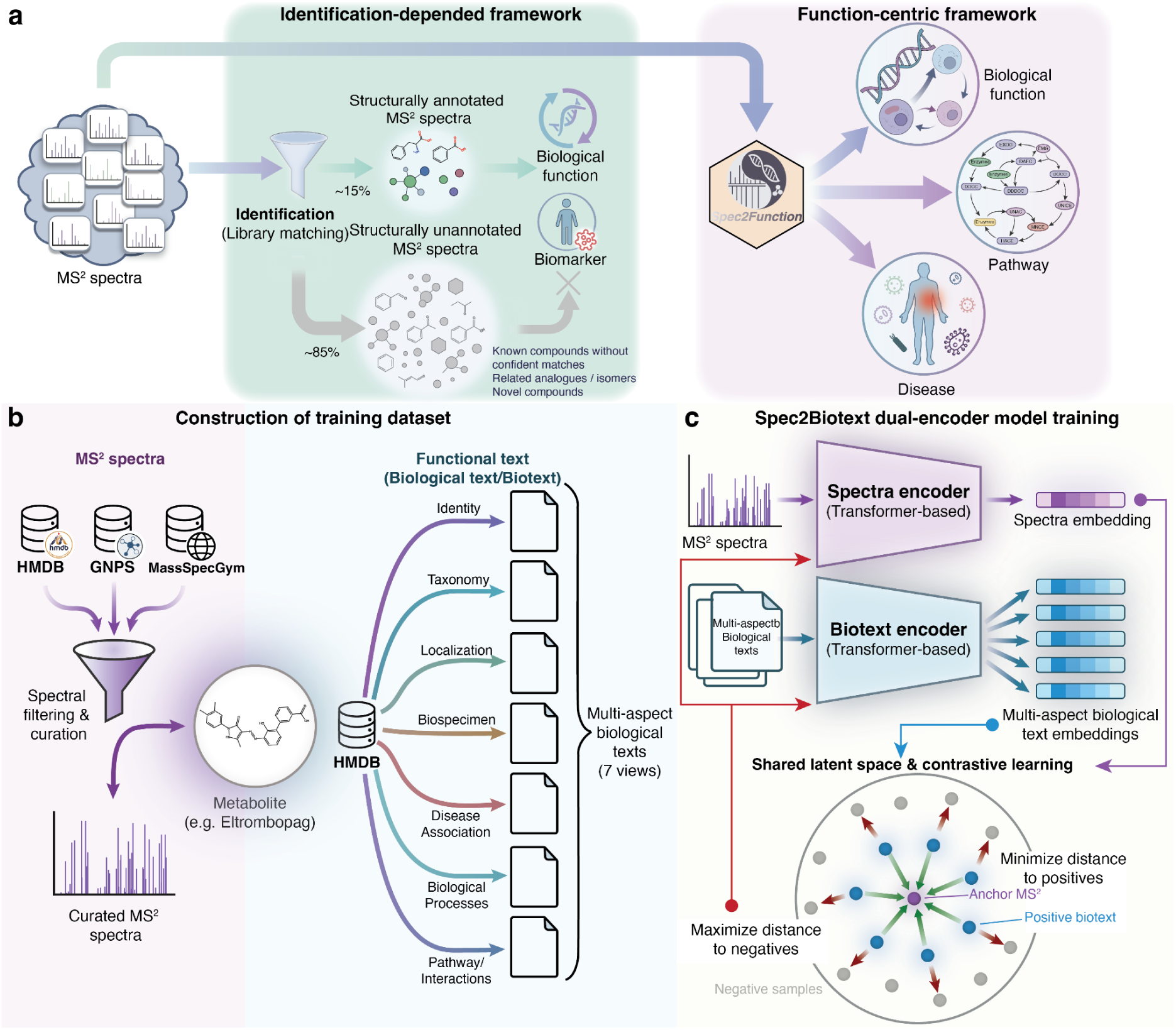
Overview of the Spec2Biotext dual-encoder model. (a) Comparison of a conventional identification-dependent workflow and the Spec2Function function-centric workflow. (b) Construction of the training dataset for Spec2Biotext. The final training dataset consisted of paired MS^2^ spectra and their corresponding biotexts. (c) Architecture and training of the Spec2Biotext dual-encoder model. The model is trained using a contrastive loss to minimize the embedding distance between matched MS^2^-biotext pairs while maximizing the distance to unmatched pairs.

To train Spec2Biotext, we constructed a paired dataset linking MS^2^ spectra to biotext annotations (Fig. 1b and Methods). MS^2^ spectra were curated from multiple public datasets and merged into a unified MS^2^ spectral database^25,38–41^. For each metabolite, we organized its function descriptions into seven aspects (Fig. 1b and Supplementary Figure 1a). This multi-aspect biotext captures both shared and distinct functional attributes among metabolites, enabling Spec2Biotext to learn shared biochemical relationships across multiple functional dimensions. Additionally, the training dataset encompassed diverse acquisition conditions and metabolite classes (Supplementary Figure 1b-f), providing broad spectral and chemical representation for model training.

With the curated training dataset in hand, we next trained our dual-encoder deep learning model to align MS^2^ spectra and biotexts in a shared representation space (Fig. 1c and Methods). For each training sample, the MS^2^ spectrum was preprocessed into tokenized *m/z* sequences paired with normalized intensity values and passed through the spectral encoder, a transformer-based neural network that outputs a fixed-length representation termed spectral embedding (Methods). In parallel, each biotext snippet associated with that metabolite was independently tokenized and encoded via a Transformer-based language model pretrained on scientific literature, producing corresponding biotext embeddings (Fig. 1c and Methods). To learn a semantically meaningful alignment between these two modalities, we adopted a contrastive learning objective^42^. The model was trained to maximize the similarity between spectral embedding and its corresponding positive biotext embeddings, while minimizing similarity to negatives (Fig. 1c). This training strategy encouraged the encoders to co-adapt and learn representations that capture shared biological semantics across multiple functional dimensions simultaneously (Supplementary Figure 2a). As a result, the spectral encoder produces spectral embeddings that implicitly encode biological function, whereas the biotext encoder learns functional embeddings that also capture structural characteristics implied by MS^2^ spectra. Once trained, the dual-encoder model enables direct comparison between spectral and biotext embeddings. Given a query MS^2^ spectrum, the model can directly identify relevant biotexts by matching query spectra against multi-aspect functional texts (Supplementary Figure 2b).

Together, Spec2Biotext aligns MS^2^ spectra with multi-aspect biological text in a unified functional space, enabling MS^2^ spectra to be directly queried for functional meaning without requiring prior structural identification of the query spectrum.

### Spec2Biotext learns a shared representation of spectral and functional information

We then evaluated whether our trained Spec2Biotext learns a biologically meaningful shared latent space in which biochemical relationships emerge through joint representation learning of MS^2^ spectra and biotext. To examine the organization of the learned biotext representation space, we encoded all metabolites using the biotext encoder and visualized the resulting embeddings using t-SNE (Fig. 2a and Methods). The biotext embeddings formed an organized functional landscape in which metabolites grouped according to broad biochemical roles. Metabolites belonging to the same chemical class, for example, sulfur-containing compounds, phosphatidylcholines, hexoses, N-acetylated sugars, and amino-acid conjugates, clustered together, indicating that the biotext embeddings of metabolites from Spec2Biotext also contain structural information (Fig. 2a).

**Fig. 2:**
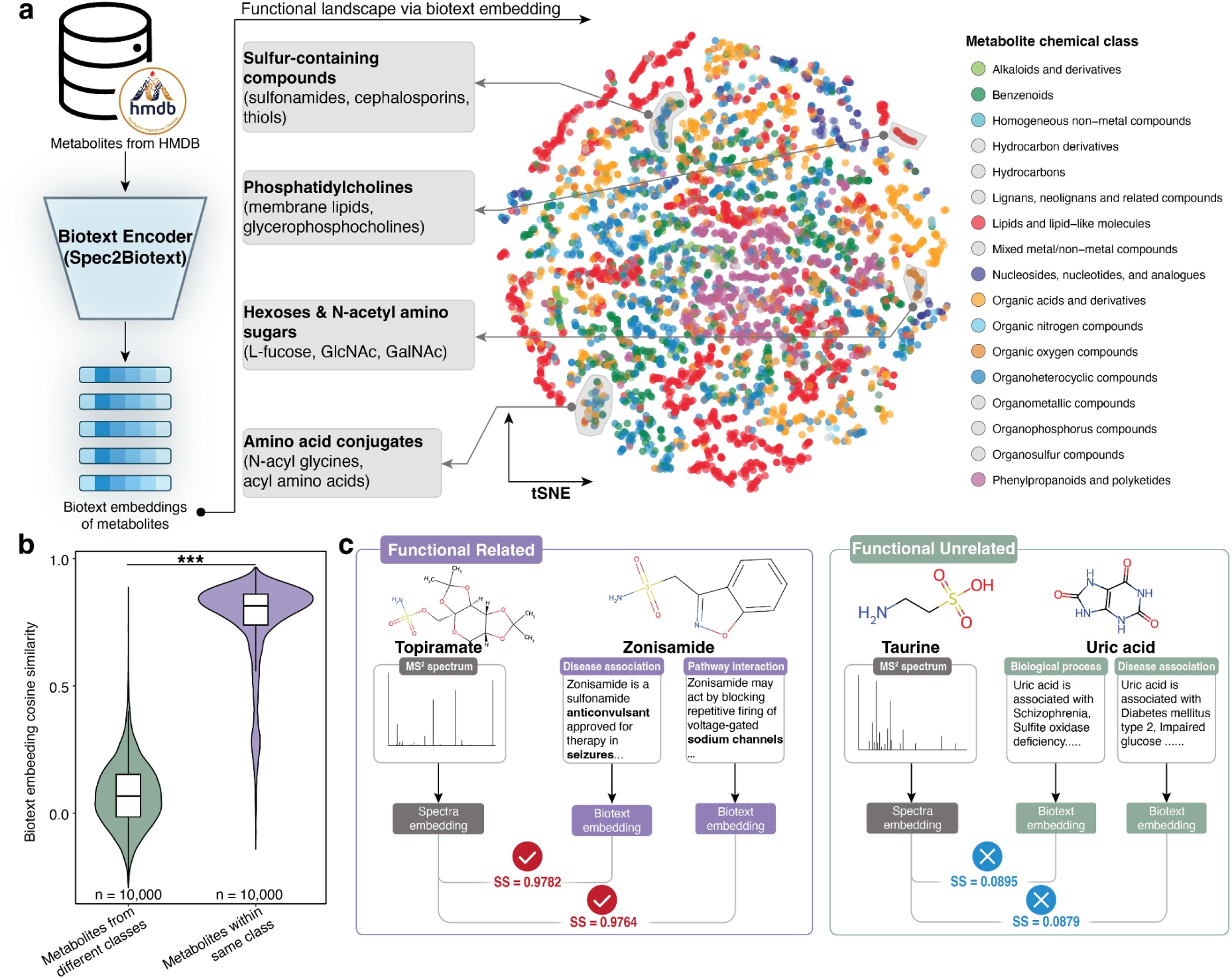
Spec2Biotext learns a cross-modal embedding space that captures functional similarity among metabolites. (a) Visualization of the functional landscape encoded by Spec2Biotext. All metabolites from the training set were encoded using the biotext encoder and projected into two dimensions using t-SNE. (b) Distribution of cosine similarities for 10,000 randomly sampled metabolite pairs from the same class compared with 10,000 randomly sampled pairs from different classes. (c) Representative functionally related and unrelated metabolite pairs illustrating metabolite functional similarity inferred by Spec2Biotext. SS: similarity score.

To quantify biochemical relatedness captured by the Spec2Biotext embeddings, we compared 10,000 randomly sampled metabolite pairs that shared both pathway membership and chemical subclass with 10,000 pairs sharing neither pathway membership nor chemical superclass (Fig. 2b and Methods). Biochemically related pairs exhibited substantially higher embedding similarity (median = 0.815, Q1-Q3: 0.739 - 0.861) than biochemically unrelated pairs (median = 0.067, Q1-Q3: -0.015 - 0.152; two-sided Wilcoxon rank-sum test, *P* < 0.001). This result further demonstrates that biotext embeddings from Spec2Biotext encode hierarchical biochemical relationships and reflect graded biochemical relatedness.

We next examined representative functionally related and unrelated metabolite pairs to illustrate the alignment between spectral embeddings and functional biotext embeddings. As an example of functionally related metabolites, we selected topiramate and zonisamide, two anticonvulsant agents with overlapping pharmacological functions despite their distinct chemical structures (Supplementary Figure 3a). We computed the cosine similarity between the spectral embedding of topiramate and two functional biotext embeddings of zonisamide (Methods). High cross-modal similarities were observed for its anticonvulsant disease-association description (0.9782) and its ion-channel- and GABA-related mechanism description (0.9764), indicating strong cross-modal functional alignment (Fig. 2c). Notably, this relationship emerged only after training the dual-encoder model. Untrained encoders failed to recover this similarity (Supplementary Figure 3b). This result demonstrates that contrastive learning is critical for aligning spectral and biological functional representations. As an example of functionally unrelated metabolites, we examined taurine and uric acid, metabolites with distinct chemical properties and biological roles (Supplementary Figure 3c). Consistent with expectation, their cross-modal embedding similarities were markedly low (biological process: 0.0895; disease association: 0.0879; Fig. 2c; Supplementary Figure 3c). Together, these representative examples illustrate cross-modal alignment patterns for functionally related and unrelated metabolites.

Taken together, these results indicate that our trained Spec2Biotext successfully integrates MS^2^ spectral and textual function information into a shared functional representation space. Contrastive learning ensures that spectral embeddings carry biologically meaningful signals, whereas biotext embeddings reflect structural regularities encoded in MS^2^ spectra. This dual-encoder model provides the conceptual and computational foundation for Spec2Function, supporting evidence-grounded biological function inference from MS^2^ spectra.

### Spec2Function: a function-centric framework enabling biological function inference directly from MS^2^ spectra

Building on the Spec2Biotext model, we developed Spec2Function, a function-centric framework for direct biological function inference from MS^2^ spectra without requiring prior structural identification of the query spectrum at inference time. To achieve this, we first constructed the Functional Embedding Metabolite Database (FemDB), a curated resource of functionally annotated metabolites embedded in a joint semantic space (Fig. 3a and Methods). Briefly, we extracted biotext descriptions for all metabolites from HMDB, encompassing diverse functional perspectives such as disease associations, pathway involvement, tissue localization, and molecular activity (Supplementary Figure 4a). These biotexts were segmented into fine-grained, semantically coherent biotext fragments and embedded using the biotext encoder from the trained Spec2Biotext model. Each resulting functional embedding is linked to the originating metabolite and its functional category, enabling semantically aware retrieval of biological context for any query MS^2^ spectrum. By indexing metabolites across multiple biological views, FemDB serves as a dense reference database for function-oriented retrieval (Fig. 3a).

**Fig. 3:**
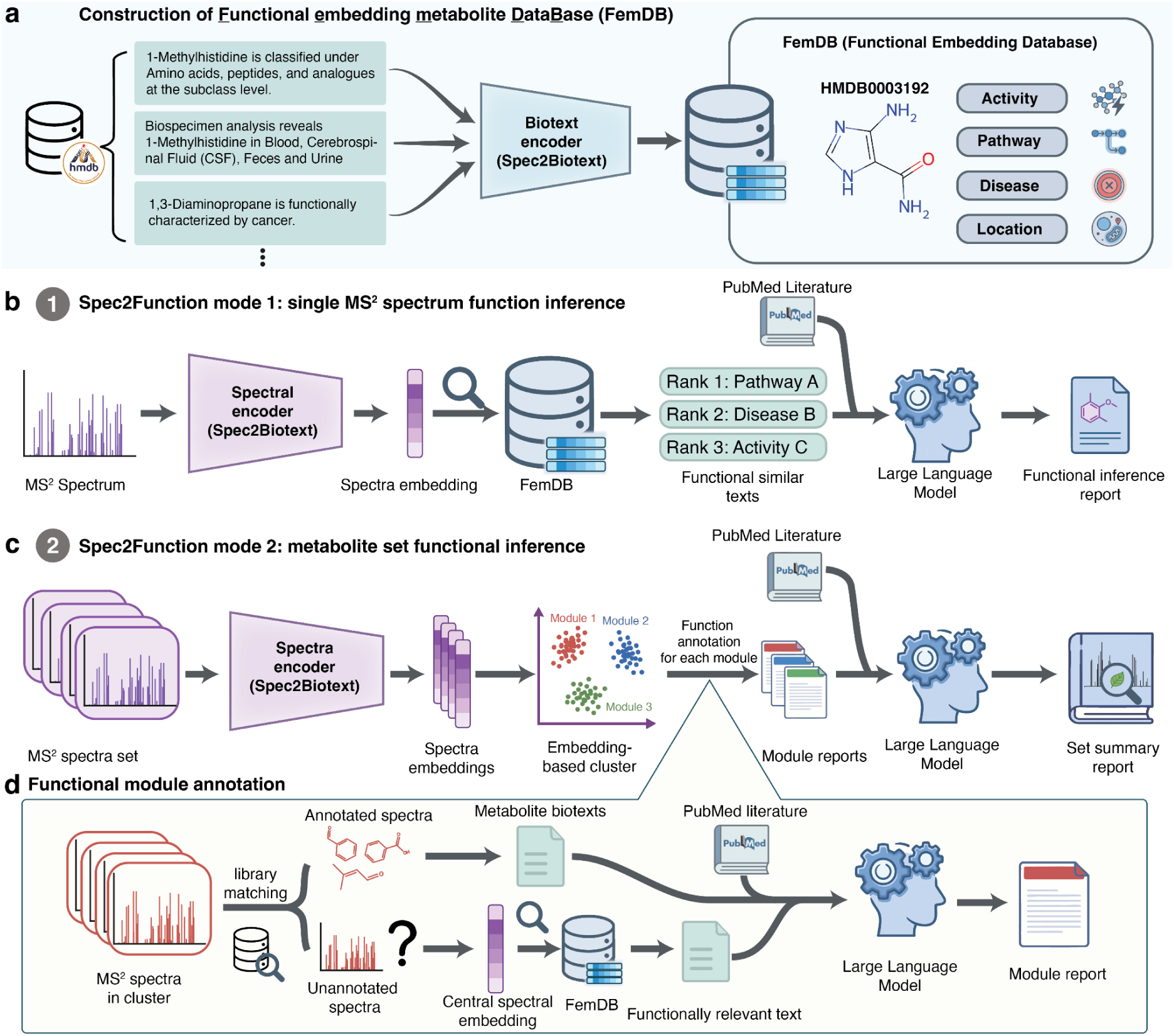
Overview of the Spec2Function framework for function-centric interpretation of MS^2^ spectra. (a) Construction of the Functional Embedding Metabolite Database (FemDB). (b) Single-spectrum function inference mode of Spec2Function. (c) Set-level function inference mode of Spec2Function. (d) Functional inference from one functional module. Within each module, unannotated MS^2^ spectra are summarized via their central spectral embedding. Functional annotation is inferred through text similarity search in FemDB, literature retrieval, and final summarization by the LLM.

Spec2Function supports two core functional inference modes tailored to different experimental designs (Fig. 3b-c and Methods). In Mode 1 (single-spectrum inference), a query MS^2^ spectrum is encoded by the spectral encoder into a latent functional embedding. This spectral embedding is used to retrieve the most similar biotext fragments from FemDB, ranked by cosine similarity (Fig. 3b). The top-ranked biotext fragments, reflecting potential biological functions of the query MS^2^ spectrum, are passed to a functional inference module that integrates external evidence from PubMed^43^. Specifically, related literature is then retrieved using metabolite names and biological terms extracted from these fragments^43^. Both the curated text fragments and the literature context are fed into an LLM using a structured, domain-specific prompt to synthesize a concise and interpretable functional summary for the spectrum (Methods and Supplementary Figure 4b). This strategy produces functional summaries grounded in curated database knowledge and supporting literature without requiring prior structural identification of the query spectrum.

In Mode 2 (set-level inference), designed for studies involving groups of MS² spectra, Spec2Function identifies functional modules from the input spectra and integrates their functional summaries into a global biological overview (Fig. 3c and Methods). First, spectral embeddings are generated for all MS^2^ spectra and subjected to unsupervised clustering in the latent space. Spectra within the same cluster are expected to share similar functional profiles, and each cluster is accordingly interpreted as a functional module. Each functional module’s centroid embedding is then used to retrieve associated biotext fragments from FemDB (Supplementary Figure 4c), and the functional inference module generates a module-level report, summarizing dominant biological themes (Fig. 3d). Finally, the functional summaries of all functional modules are integrated to generate a global overview of the entire MS^2^ spectral landscape (Fig. 3c).

Taken together, Spec2Function provides a function-centric framework for LC-MS/MS untargeted metabolomics that supports functional inference beyond metabolites represented during model development. By leveraging the representational power of dual encoders, a densely annotated multimodal database, and literature-aware language modeling, Spec2Function can generate evidence-grounded functional hypotheses for individual MS^2^ spectra and spectrum sets without requiring prior structural identification of the query spectra at inference time.

### Spec2Function supports functional inference across metabolite- and pathway-level benchmarks

To assess the performance of Spec2Function, we first evaluated its ability to infer the biological functions of individual metabolites using an external MS^2^ spectral dataset (Fig. 4a and Methods). This validation set consisted of 404 metabolites, each with an experimentally acquired MS^2^ spectrum and a matched HMDB ID, serving as ground-truth annotations (Supplementary Data 1).

**Fig. 4:**
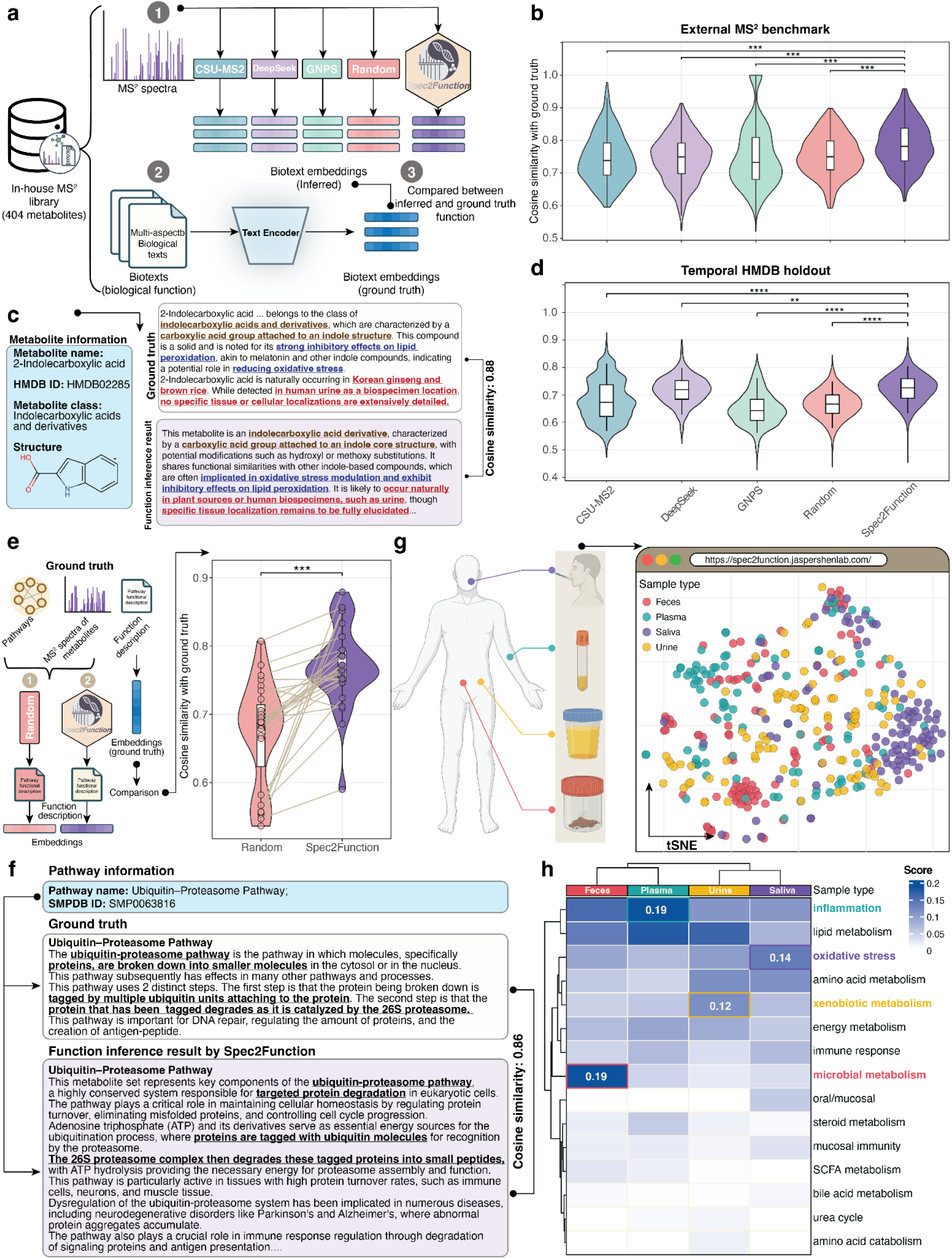
Quantitative benchmarking, temporal generalization, and biofluid-specific functional profiling of Spec2Function. (a) Overview of the Spec2Function benchmarking strategy. (b) Violin plots showing cosine similarity between inferred and ground-truth biotexts across different methods in the single-metabolite benchmark. (c) Representative example from the single-metabolite benchmark using 2-indolecarboxylic acid. (d) Temporal HMDB holdout benchmark using metabolites introduced in HMDB v5 but absent from the frozen HMDB v4 training and retrieval resources. Violin plots compare functional inference performance across different methods. (e) Pathway-level benchmark of Spec2Function using 27 metabolic pathways. Violin plots compare the cosine similarity of inferred versus ground-truth functions for Spec2Function and the spectrum-shuffled random control. (f) Representative example from the pathway-level benchmark using the Ubiquitin-Proteasome Pathway. (g) Application of Spec2Function to functional profiling of four human biofluids: feces, plasma, urine, and saliva. (h) Heatmap showing the proportion of metabolites associated with each functional category across four biological sample types. Rows represent functional categories derived from keyword analysis of Spec2Function-inferred annotations; columns represent sample types (feces, saliva, plasma, and urine). Color intensity reflects the proportion of metabolites annotated with each functional category within each sample type.

For each metabolite, we queried Spec2Function to retrieve biological functions based on the MS^2^ spectrum alone. These inferred descriptions were compared with functional descriptions (ground truth) using a text-based cosine similarity metric. We benchmarked Spec2Function against four alternative strategies (CSU-MS2^33^, DeepSeek-V3^44,45^, GNPS^40^, and Random), which span spectrum-to-structure retrieval, direct LLM inference, spectral-library matching, and retrieval ablation (Methods). Spec2Function achieved the highest median similarity between predicted and ground truth functional descriptions, significantly outperforming all comparator methods (Wilcoxon signed-rank test, all *P* < 0.001; Fig. 4b).

As an illustrative zero-shot example, we examined 2-indolecarboxylic acid (Fig. 4c). Spec2Function recovered several key attributes of this metabolite, including its structural class (indolecarboxylic acids), functional roles (oxidative stress modulation and lipid metabolism inhibition), and occurrence in biological specimens and natural sources (*e.g.*, urine and ginseng) (Fig. 4c). This example illustrates that Spec2Function can recover multiple functional attributes from an MS^2^ spectrum.

To more stringently evaluate generalization to metabolites absent from both model training and the retrieval database, we performed a temporal HMDB holdout analysis (Fig. 4d and Methods). Spec2Biotext and FemDB were reconstructed and frozen using HMDB v4, whereas metabolites newly introduced in HMDB v5 and absent from the frozen v4 knowledge base were used for testing. The final benchmark comprised 673 temporally held-out metabolites represented by 7,153 positive-ion MS^2^ spectra. Spec2Function achieved the highest mean similarity to the corresponding v5 reference biotexts (0.723), compared with DeepSeek-V3 (0.717), CSU-MS2 (0.679), Random (0.666), and GNPS (0.649). The improvement over DeepSeek-V3 was significant (paired Wilcoxon test, *P* = 0.0071), with larger differences observed relative to CSU-MS2, GNPS, and Random (all *P* < 0.001). These results demonstrate that Spec2Function can transfer learned spectrum–function relationships to metabolites absent from both model training and the frozen retrieval database.

To reduce unsupported functional assignments, Spec2Function applies a cosine-similarity-based confidence gate and abstains from prediction when retrieval support is insufficient. On the external benchmark, below-threshold queries showed significantly lower annotation quality when predictions were forcibly generated, supporting the effectiveness of this confidence-aware abstention strategy (Supplementary Note and Supplementary Figure 6).

We further examined the robustness of Spec2Function across different mass spectrometry platforms using matched QTOF and Orbitrap spectra from 2,728 metabolites. Spec2Function was trained using spectra acquired on one instrument platform and evaluated on matched spectra from the same metabolites acquired on the other platform, and vice versa. The model maintained consistent functional inference performance across the two instrument settings, supporting cross-platform transfer of the learned spectrum-function representation under the tested conditions (Supplementary Figure 6b-c).

We next evaluated the ability of Spec2Function to infer the biological functions of entire metabolic pathways (Methods). Twenty-seven pathways with experimentally validated MS^2^ spectra for their constituent metabolites were selected (Fig. 4e and Supplementary Data 2). MS^2^ spectra from each pathway were aggregated and input into Spec2Function, which generated pathway-level functional annotations by integrating information from all associated spectral embeddings. A random control was constructed by shuffling spectra across pathways. Functional similarity to the ground-truth annotations was again assessed using cosine similarity (Fig. 4e). Spec2Function achieved significantly higher similarity scores than the random control (median similarity: 0.7652; IQR: 0.7495-0.7939), indicating strong functional coherence (Fig. 4e). In contrast, the random baseline exhibited poor similarity (median: 0.6878; IQR: 0.6233-0.7141), validating the specificity of Spec2Function’s predictions (Wilcoxon signed-rank test, *P* < 0.001). As a zero-shot case study, we highlight the Ubiquitin-Proteasome Pathway (Fig. 4f), where Spec2Function not only recapitulated the exact pathway name but also correctly captured the core biological functions, including proteasomal protein degradation, immunological signaling, and disease relevance in neurodegeneration (Fig. 4f).

Finally, as an application-level demonstration, we applied Spec2Function to LC-MS/MS untargeted metabolomics datasets from four distinct human biofluids (feces, plasma, urine, and saliva) (Fig. 4g and Methods). For each dataset, all available MS^2^ spectra were processed using Spec2Function, and the distribution of inferred biological functions was summarized (Fig. 4g). The resulting functional profiles showed strong alignment with known tissue physiology (Fig. 4h). Specifically, fecal samples were enriched in microbial metabolism functions, reflecting their direct contact with gut microbiota^46^. Plasma samples showed dominant signals related to inflammation, consistent with the role of circulating immune mediators^47^. Urine samples exhibited the highest enrichment for xenobiotic metabolism, in line with renal excretion of exogenous compounds^48,49^. Strikingly, saliva samples were uniquely enriched for oxidative stress functions, an observation supported by prior studies linking salivary redox imbalance to oral and systemic diseases^50^.

Additionally, a paired comparison with matched reference spectra further showed that functional annotation performance was largely preserved for spectra acquired directly from biological samples, with no significant difference detected under the tested conditions (Supplementary Figure 6 and Methods). Together, these results indicate that Spec2Function enables accurate and robust functional inference for both individual metabolites and entire metabolic pathways across diverse contexts, and that its RAG design reduces erroneous or hallucinated annotations (Supplementary Note and Supplementary Figure 5).

### Spec2Function expands functional coverage in liver-cirrhosis metabolomics

To assess the breadth and depth of biological insights enabled by Spec2Function, we conducted a systematic comparison between Spec2Function and an identification-dependent workflow (Methods). As a representative test case, we analyzed a well-characterized plasma metabolomics study of liver cirrhosis (Fig. 5a and Methods)^51^. Plasma samples were profiled using LC-MS/MS, and the raw data in positive-ion mode were processed using Spec2Function and, in parallel, an identification-dependent workflow implemented using the TidyMass ecosystem^7,8^. Differential analysis identified 442 peaks with MS^2^ spectra that were significantly altered between two groups (182 upregulated and 260 downregulated; Fig. 5b). Consistent with the persistent limitations of spectral libraries, only 113 of these peaks (25.6%) could be structurally identified, indicating that approximately three-quarters of these differential features could not enter this identification-dependent pathway analysis (Fig. 5c).

**Fig. 5:**
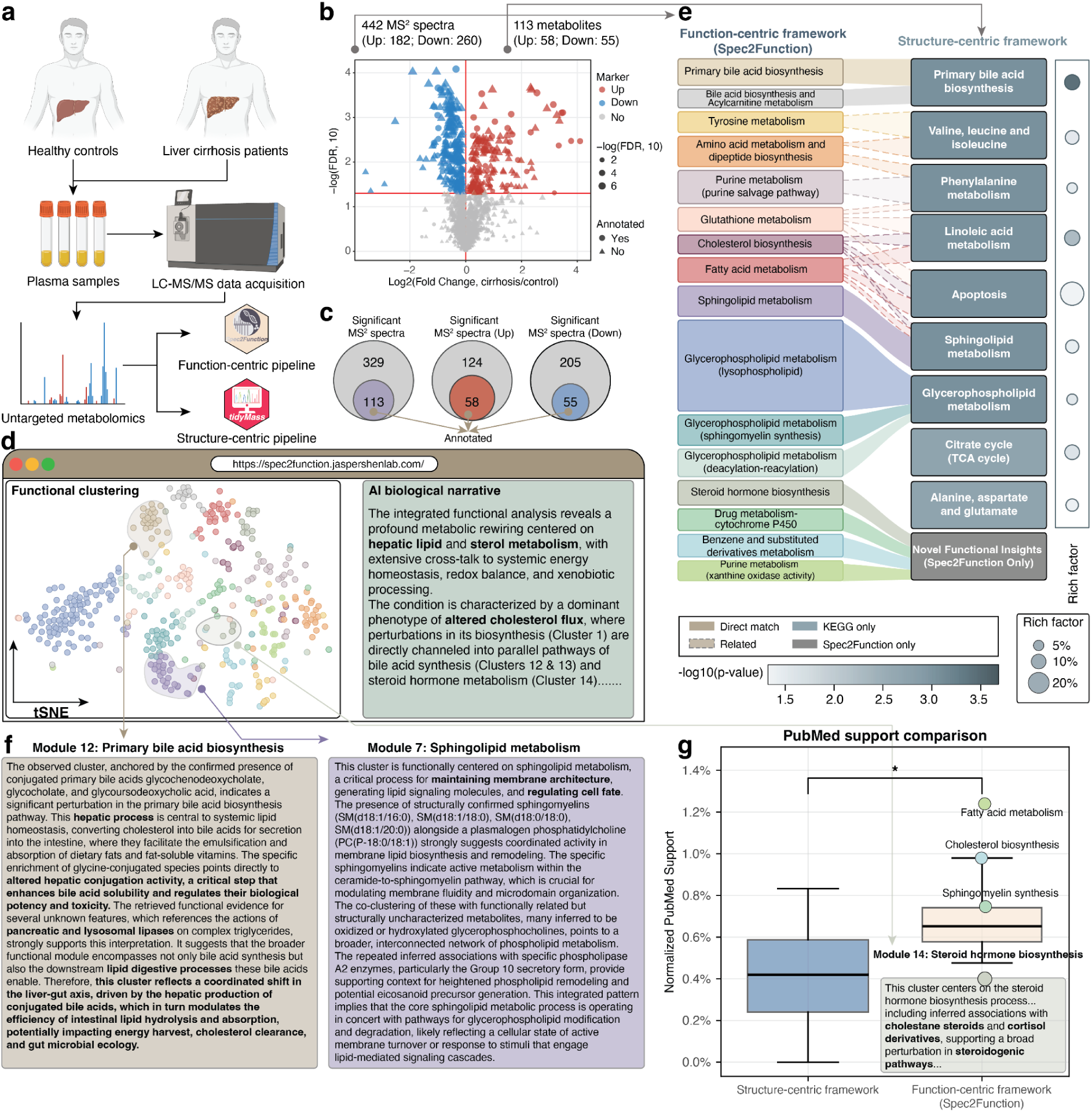
Spec2Function enables more comprehensive metabolic functional discovery. (a) Overview of the liver cirrhosis plasma metabolomics dataset analyzed using both the identification-dependent workflow and Spec2Function. (b) Volcano plot illustrating 442 differential peaks (with MS^2^ spectra) between cirrhosis and healthy controls (positive mode). (c) Identification rates of significant MS^2^ spectra. (d) Functional landscape inferred by Spec2Function. (e) Comparative mapping between Spec2Function-derived functional modules and enriched pathways obtained from the identification-dependent workflow. (f) Representative Spec2Function-derived functional module interpretations, showing accurate and mechanistically grounded descriptions of liver-associated processes. (g) Leakage-controlled and frequency-normalized PubMed literature support for functional modules and pathways uniquely identified by either Spec2Function or the identification-dependent workflow.

Applying Spec2Function to the full set of 442 differential MS^2^ spectra produced a broad functional landscape (Fig. 5d). Without requiring prior structural identification of every input feature, Spec2Function recovered hallmark metabolic perturbations associated with liver cirrhosis, most notably disruptions in hepatic lipid metabolism, sterol metabolism, and cholesterol processing, each of which is well established in cirrhotic pathology^52^. Additionally, 16 coherent functional modules spanning diverse biological processes were identified (Fig. 5d; Supplementary Data 3). These included bile acid synthesis, sphingolipid remodeling, amino-acid metabolism, xenobiotic detoxification, and oxidative stress regulation (Supplementary Note).

By contrast, applying the identification-dependent workflow to the same dataset yielded nine significantly enriched metabolic pathways derived from the 113 identifiable metabolites (Fig. 5e; Supplementary Data 4). To compare results between two frameworks, we assessed functional concordance between Spec2Function-derived functional modules and pathways from the identification-dependent workflow (Supplementary Figure 7a). Several processes, most notably primary bile acid biosynthesis and sphingolipid metabolism, were consistently recovered by both approaches (Fig. 5e), reflecting their well-established roles in hepatic dysfunction^53,54^. Spec2Function, however, provided a broader context for these modules by integrating cross-modal evidence, including literature-supported text fragments (Fig. 5f and Supplementary Note).

To further assess whether these findings depended on the choice of comparator, we benchmarked the same liver-cirrhosis dataset using two additional metabolite-level annotation workflows: CSI:FingerID^55^, representing molecular-formula- and candidate-structure-based annotation, and MetDNA^56^, representing spectral-similarity- and reaction-network-based annotation propagation (Supplementary Figure 8 and Methods). CSI:FingerID generated top-ranked structural candidates for 370 of the 442 differential features and, after restricting pathway analysis to the highest-confidence predictions, recovered canonical liver-associated pathways including primary bile acid biosynthesis and cholesterol metabolism. MetDNA matched most differential features to reconstructed features but yielded KEGG annotations for only 15 features; nevertheless, these annotations recovered a strong fatty acid degradation signal. Thus, these independent structure-assisted workflows supported several of the major lipid, bile-acid, and sterol perturbations recovered by Spec2Function, while their downstream pathway analyses remained dependent on the subset of features that could be assigned usable metabolite identities (Supplementary Note).

MS2MP^19^, in contrast, generated compatible outputs for 389 of the 442 differential features, of which 278 received at least one pathway label (Supplementary Note and Supplementary Figure 8). However, its leading enriched pathways were dominated by non-ribosomal peptide, macrolide, polyketide, and xenobiotic degradation terms that were less specific to human liver-cirrhosis biology. Consistent with this observation, PubMed-based disease-relevance analysis showed significantly stronger liver-cirrhosis support for Spec2Function-derived functional outputs than for MS2MP predictions (*P* = 0.0481; Supplementary Figure 8). Together, these comparisons indicate that Spec2Function combines broad utilization of structurally unannotated MS^2^ features with disease-relevant functional specificity, bridging the trade-off between annotation coverage and biological interpretability observed across the alternative workflows.

Notably, four functional modules uniquely identified by Spec2Function were absent in the structure-centric results. Inspection revealed that these modules were driven largely by MS^2^ spectra lacking structural assignments (Supplementary Figure 7b), explaining why structure-centric processing could not detect them. To assess phenotype-related literature support for these modules while minimizing evidence overlap and publication-frequency bias, we performed a leakage-controlled and frequency-normalized PubMed analysis using NCBI E-utilities, excluding publications used during Spec2Function module generation and quantifying literature support as the proportion of publications associated with each functional term that also co-occurred with “liver cirrhosis” (Supplementary Figure 7c and Methods). The most strongly supported Spec2Function modules included fatty acid metabolism (1.24%), cholesterol biosynthesis (0.98%), and glycerophospholipid remodeling (0.73-0.75%), collectively highlighting broad lipid-metabolic dysregulation in liver cirrhosis (Fig. 5g). Additional Spec2Function-specific modules, including steroid hormone biosynthesis, purine metabolism, and cytochrome P450-mediated drug metabolism, were also supported by independent literature evidence after leakage control (Supplementary Note). For example, dysregulated steroid hormone metabolism is well documented in cirrhosis but was not recovered by the identification-dependent workflow.

Conversely, two pathways identified only by the identification-dependent workflow, citrate cycle (TCA cycle) and alanine, aspartate and glutamate metabolism, showed no literature support for relevance to cirrhosis and were based on sparse metabolite coverage (2 of 20 and 2 of 28 metabolites mapped, respectively). We therefore interpret these enrichments cautiously, particularly given their sparse metabolite coverage (Supplementary Figure 7d).

Collectively, these comparisons show that Spec2Function extends functional analysis to a substantially larger fraction of disease-associated MS^2^ features while retaining biologically coherent and disease-relevant interpretations. Structure-assisted approaches independently recovered several canonical metabolic alterations also identified by Spec2Function, particularly bile acid, cholesterol, and fatty-acid metabolism, but their downstream pathway analyses remained dependent on metabolite-level annotation. By contrast, MS2MP achieved broad spectral coverage but showed lower disease specificity among its leading pathway predictions. Spec2Function therefore provides a complementary function-centric strategy that combines broad MS^2^ feature utilization with biologically specific interpretation, enabling disease-associated metabolic signals to be analyzed even when confident structural assignments are unavailable.

### Spec2Function reveals comprehensive metabolic alterations across organs during macaque aging

To evaluate Spec2Function in a biological context, we applied it to a previously generated multi-organ metabolomics dataset from female macaques (*Macaca mulatta*) (Fig. 6a)^57^. Across tissues, the number of age-associated metabolic peaks ranged from 3 to 1057, with a median of 442 peaks per tissue, prior to structural annotation. As is common in untargeted metabolomics datasets, only a small fraction of these peaks could be structurally annotated. Using the TidyMass-based identification-dependent workflow (Methods), the identification rate of age-associated peaks ranged from 0% to 17.9% (median 9%) across the 30 tissues (Fig. 6a), leaving the majority of differential MS^2^ spectra without confident structural annotations.

**Fig. 6:**
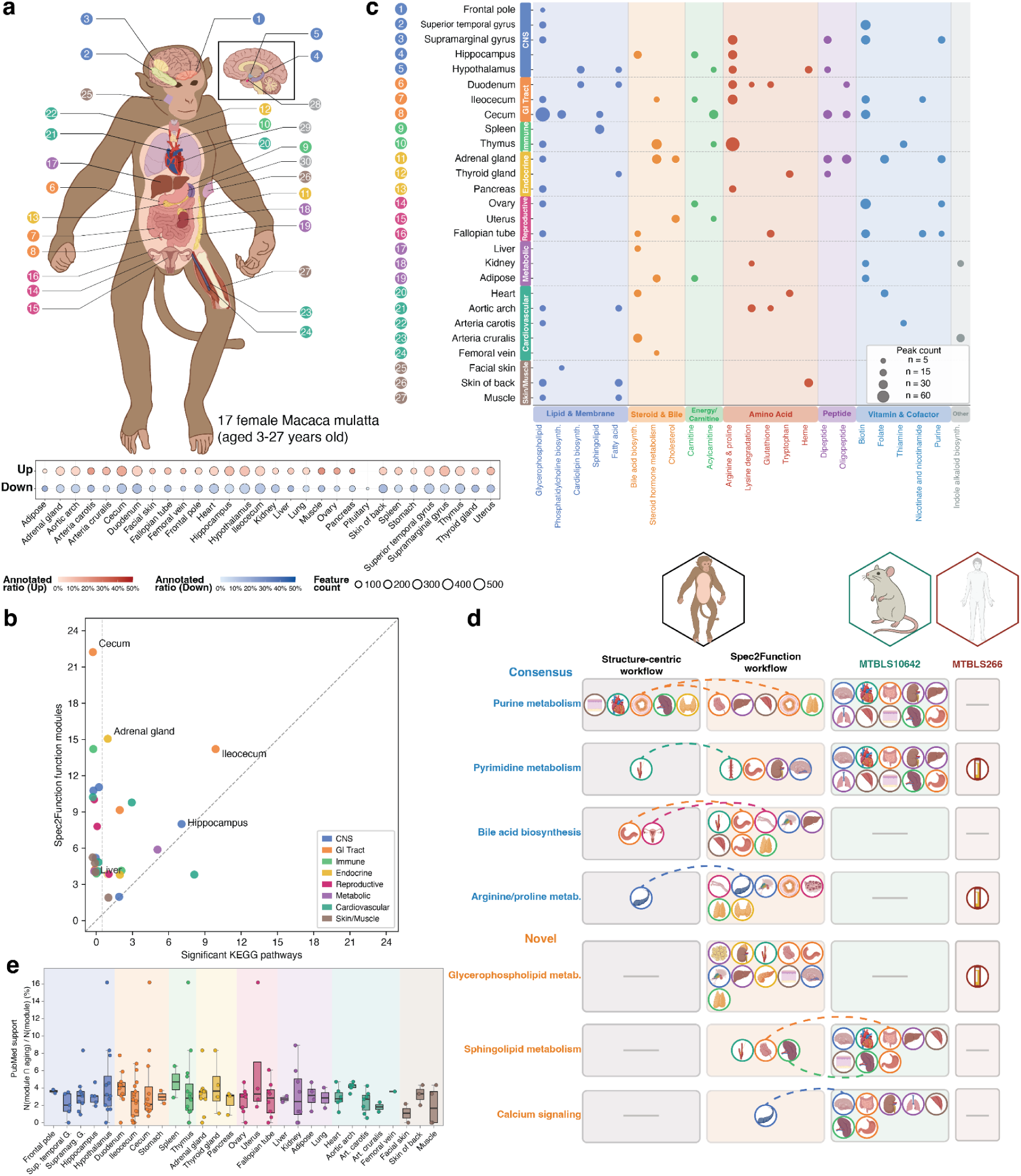
Spec2Function reveals comprehensive metabolic functional alterations across organs during aging. (a) Overview of the multi-organ macaque aging metabolomics dataset. The lower panel shows the proportion of annotated metabolites among significantly upregulated (red) and downregulated (blue) peaks across tissues. (b) Comparison between the number of functional modules identified by Spec2Function and the number of pathways detected using an identification-dependent workflow. (c) Tissue distribution of metabolic functional modules inferred by Spec2Function across organs. Functional modules detected in at least two tissues are shown. Circle size indicates the number of MS^2^ spectra contributing to the inferred function. (d) Comparison between pathways identified by the identification-dependent workflow and functional modules inferred by Spec2Function. (e) Publication-excluded, frequency-normalized aging-related literature support for Spec2Function-derived functional modules.

We next applied Spec2Function to all differential MS^2^ spectra across the 30 tissues. In parallel, we analyzed the same dataset using an identification-dependent workflow in which pathway enrichment analysis was performed on all successfully annotated metabolites (Methods). Across tissues, Spec2Function identified more functional modules compared with the identification-dependent workflow (Fig. 6b). For visualization, we focused on functional modules detected in at least two tissues (Fig. 6c).

The inferred modules showed marked tissue specificity and recapitulated metabolic programs associated with major physiological systems (Fig. 6c). Glycerophospholipid metabolism was the most prevalent module and was detected in 14 tissues. This distribution is consistent with the broad roles of glycerophospholipids in membrane organization, membrane dynamics, intracellular signaling, and organelle function^58,59^, as well as with prior reports of age-associated lipid remodeling across tissues^60,61^. Arginine and proline metabolism was the second most prevalent module and was detected in seven tissues, with the strongest signal observed in the thymus. This pattern is consistent with established links between arginine metabolism and immune regulation during aging^62,63^, and between proline metabolism, mitochondrial energetics, and redox homeostasis^64,65^.

Several tissue-enriched modules were also identified (Fig. 6c). Steroid hormone metabolism was most prominent in the adrenal gland, adipose tissue, and thymus, consistent with known roles of these tissues in steroid production, interconversion, or responsiveness^66,67^. Bile acid metabolism was detected in the liver, as expected, and was also observed in extrahepatic tissues, including the hippocampus and heart. Although these tissues are not canonical sites of bile acid biosynthesis, bile acids have been increasingly recognized as systemic signaling molecules involved in metabolic and inflammatory regulation across organs^54,68^. In cardiovascular tissues, bile acid signaling through receptors such as FXR and TGR5 has been implicated in metabolic and inflammatory control^69,70^.

We next compared the inferred modules with pathways identified by the identification-dependent workflow (Methods). Modules were classified into two groups: consensus pathways, detected by both workflows in the macaque dataset, and Spec2Function-specific modules, detected only by Spec2Function under the analysis settings used here. Several established aging-associated metabolic programs were recovered by both approaches (Fig. 6d), including purine metabolism, pyrimidine metabolism, bile acid metabolism, and arginine and proline metabolism. In several cases, both workflows converged on the same tissues, including bile acid metabolism in the duodenum and arginine and proline metabolism in the hippocampus^71,72^. These results indicate that Spec2Function recapitulates metabolic signals also captured by the identification-dependent pathway analysis.

Spec2Function also identified additional modules that were not detected in the identification-dependent workflow (Fig. 6d). These included glycerophospholipid metabolism, sphingolipid metabolism, and calcium signaling, which were not reported as prominent aging-associated programs in the original macaque study^57^. Glycerophospholipid metabolism was detected in 11 tissues and showed concordant enrichment in an independent human aging plasma dataset (Hypergeometric test, *P* < 0.001). Sphingolipid metabolism was also enriched in matched mouse organs from an independent aging atlas dataset, including spleen and intestine, corresponding to tissues in which Spec2Function identified the same functional theme in macaques. Similarly, calcium signaling, detected by Spec2Function in the macaque hippocampus, was enriched in the aging mouse brain (Hypergeometric test, *P* = 0.03).

To further assess phenotype-related literature support while minimizing evidence overlap and publication-frequency bias, we applied the same analysis used in the liver-cirrhosis study (Methods). The resulting normalized support values are shown in Fig. 6e and are interpreted as phenotype-related literature support rather than independent evidence of biological correctness. Together, these results show that Spec2Function recapitulates established aging-associated metabolic programs while also identifying additional functional modules not recovered by the identification-dependent analysis. The concordance observed across macaque, mouse, and human datasets further supports the relevance of these inferred modules to aging biology.

## Discussion

LC-MS/MS untargeted metabolomics has long been limited by its reliance on structural identification, which remains challenging because of incomplete spectral libraries, instrument-dependent variation, and limited structural annotation coverage. As a result, only a minority of detected peaks in a typical dataset can be assigned a confident structure, leaving a substantial proportion of metabolomics signals difficult to interpret biologically. Here we present Spec2Function, a framework for function-oriented interpretation of MS^2^ spectra. By separating query-time functional inference from prior structural identification of the query spectrum, Spec2Function expands the fraction of untargeted metabolomics data that can be incorporated into downstream biological analysis. This distinction does not imply independence from characterized biological knowledge during model development; rather, it separates the knowledge used to train and construct the retrieval resources from the information required for a new query at inference time.

A central component of this framework is Spec2Biotext, a dual-encoder model that maps MS^2^ spectra and multi-aspect metabolite biotext into a shared embedding space. Through contrastive learning on spectrum-text pairs, the model learns associations between fragmentation patterns and semantic descriptions of metabolite-related properties and functions. In our analyses, this embedding space captured biochemical organization across multiple levels, from broad chemical classes to cross-metabolite functional relationships, supporting the use of shared spectral-text embeddings for function-oriented retrieval.

Building on this foundation, Spec2Function provides evidence-grounded functional inference for individual spectra and sets of differential features. Across the revised benchmarks, Spec2Function showed higher functional-description similarity than the evaluated comparators and retained measurable performance in the temporal HMDB holdout, while the pathway-level benchmark demonstrated coherent functional recovery. The retrieval-augmented generation module grounds downstream summarization in FemDB-derived evidence and related literature, reducing unsupported functional assignments relative to direct LLM prompting in our benchmarks. Together with confidence-aware abstention, this design provides a function-centric strategy for evidence-grounded functional summarization.

The liver-cirrhosis analysis illustrates how Spec2Function complements existing metabolomics workflows. The original spectral-library-based analysis was expanded to include SIRIUS/CSI:FingerID, MetDNA, and MS2MP, representing candidate-structure annotation, annotation propagation, and direct MS^2^-to-pathway inference, respectively. These approaches recovered overlapping but distinct aspects of liver-cirrhosis metabolism. Structure-assisted workflows recovered canonical bile-acid, cholesterol, and fatty-acid signals when usable metabolite-level annotations were available, whereas MS2MP provided broad pathway-level coverage. Spec2Function used the full differential MS^2^ feature set to construct evidence-grounded functional modules, including contributions from structurally unannotated spectra. These comparisons emphasize complementary coverage and output types rather than superiority of one broad analytical paradigm over another.

By applying Spec2Function to a multi-organ macaque aging dataset, we demonstrate the framework’s utility for biological discovery. Spec2Function reconstructed the metabolic functional landscape across 30 organs and revealed both organ-specific functional rewiring and shared hallmarks of aging. Moreover, Spec2Function identified several aging-associated functional modules absent from the identification-dependent workflow, which were subsequently validated across independent human and mouse datasets. These results suggest that a function-centric approach can reveal biology that is not recovered by the identification-dependent workflow. The cross-species concordance with independent human and mouse metabolomics datasets provides a separate line of evidence that is independent of the PubMed literature-support analysis.

Despite these advances, Spec2Function has limitations. First, the functional embedding and retrieval space is constrained by the coverage and quality of curated biotext and FemDB; understudied metabolites and chemical regions distant from the represented knowledge space may have insufficient support. Second, the current implementation focuses on positive-ion MS^2^ spectra and is primarily human-centric because its functional knowledge base is derived largely from HMDB. We therefore do not assume equivalent performance for species-specific metabolomes or distant taxonomic groups. Third, spectral quality remains important: severe loss or distortion of informative fragments can reduce retrieval support and lead to abstention. Fourth, the final LLM-generated interpretation depends on the quality of the retrieved evidence and should be regarded as an evidence-grounded functional hypothesis rather than a definitive mechanistic assignment. Future work may expand FemDB with additional pathway and cross-species resources and integrate complementary structural-prediction methods to broaden applicability.

Overall, Spec2Function provides a complementary route for interpreting MS^2^ spectra in untargeted metabolomics. Rather than replacing existing metabolite identification workflows, it is intended to work alongside them: function-centric analysis can be used to prioritize biologically relevant signals, after which key metabolites can still be subjected to targeted structural characterization using established analytical approaches. In this way, Spec2Function offers a practical strategy for extracting biological value from the large fraction of untargeted metabolomics data that remains structurally unannotated.

## Methods

### Model architecture, training strategy, and data processing

Spec2Biotext adopts a dual-tower contrastive learning architecture to learn semantic alignments between MS^2^ spectra and biomedical text descriptions. One encoder processes MS^2^ spectra (spectral encoder), and another encodes biomedical text (biotext encoder), with both projecting their outputs into a shared embedding space to support cross-modal retrieval and annotation. To improve the contextual representation of biological text, we incorporated an auxiliary masked language modeling (MLM) objective on the biotext encoder^73^.

The model was trained on MS^2^-biotext pairs derived from the Human Metabolome Database (HMDB), where each biotext is a structured natural language description generated from standardized templates applied to HMDB’s annotated fields. After training, Spec2Biotext can embed an input MS^2^ spectrum satisfying the supported preprocessing and *m/z* requirements into the shared semantic space and retrieve functionally relevant biotext representations from FemDB, a functional embedding database constructed from HMDB-derived biotexts. Characterized structural and biological knowledge is required to construct the training resources, whereas prior structural identification of a new query spectrum is not required for inference. Generalization beyond the development resources is evaluated explicitly using the temporal HMDB holdout benchmark described below.

### MS^2^ spectral database

We curated MS^2^ spectra from multiple public databases, including the Human Metabolome Database (HMDB)^25^, MassBank of North America (MoNA)^38^, MassBank, the Global Natural Products Social Molecular Networking platform (GNPS)^40^, and MassSpecGym^41^ (Supplementary Data 5). Only the spectra associated with valid HMDB compound identifiers were retained for downstream analysis. After filtering, we obtained a total of 8,438 unique metabolites corresponding to 193,755 MS^2^ spectra.

The dataset was further stratified by ionization mode. In positive mode, the final set contained 8,149 metabolites and 174,841 MS^2^ spectra, while in negative mode, the final set contained 2,132 metabolites and 18,715 MS^2^ spectra. Due to the comparatively limited coverage in negative mode, we focused exclusively on the positive-ion mode spectra for subsequent model training and evaluation. Each MS^2^ spectrum was normalized such that the most intense fragment within the spectrum was scaled to a relative intensity of 1.0, preserving the relative peak intensities for model input.

### Training dataset

The Human Metabolome Database (HMDB) was downloaded from https://www.hmdb.ca/ on March 18, 2025. The full release contained 248,138 metabolite entries, each annotated with rich biological and chemical information, including functional summary, metabolic pathways, biospecimen and tissue localizations, disease and protein associations, and standard chemical identifiers (*e.g.*, InChIKey, SMILES, PubChem ID).

#### Biotext generation

For each metabolite in HMDB, we systematically extracted and standardized all available biological and chemical annotations from the structured XML records to construct biological function texts, referred to as biotexts. Rather than employing a large language model, we designed standardized natural language templates for each functional field in the HMDB XML schema, converting structured database entries into model-readable text fragments. Seven functional categories were covered: (1) molecular identity, including compound names and chemical classifications; (2) taxonomic hierarchy, encompassing kingdom, superclass, class, subclass, and direct parent levels along with chemical descriptions; (3) biological localization, including cellular compartments and tissue localizations; (4) biospecimen distribution, covering the presence of metabolites across different biological specimens; (5) pathway and protein interactions, covering metabolic pathway memberships and associated proteins such as enzymes and transporters; (6) disease associations, capturing known links between metabolites and human diseases; and (7) biological processes, including ontology-derived functional terms describing biological roles and processes. All templates used for biotext generation are provided in Supplementary Data 6.

To increase textual diversity, multiple sentence templates were defined for each category and randomly sampled during generation. In total, 2,515,806 biotext records were generated from 248,138 HMDB metabolite entries, spanning Taxonomy (1,281,867), Biological Processes (463,414), Pathway/Interactions (413,010), Identity (245,324), Localization (45,673), Biospecimen (42,776), and Disease Association (23,742). For visualization, biotext embeddings were projected into two dimensions using t-SNE (scikit-learn implementation) with perplexity = 70, max iterations = 1,000, PCA initialization, and automatic learning rate.

#### Functional description for evaluation

To evaluate the quality of generated functional annotations, we constructed a separate evaluation corpus consisting of LLM-generated functional descriptions for all HMDB metabolites. Each summary was produced by prompting a large language model (GPT-4o-mini-2024-07-18) with the full structured HMDB record for a given metabolite, instructing it to generate a concise, coherent 3–6 sentence paragraph covering the metabolite’s chemical classification, biological localizations, pathway involvement, and known physiological functions (Supplementary Note). These LLM-generated functional descriptions served exclusively as ground-truth references for evaluating functional annotation quality.

#### Training sample pairs

Because a single metabolite may be associated with multiple MS^2^ spectra, these were all included as independent training examples to enhance spectral diversity. After combining all MS^2^ spectra per metabolite with the available biotexts, the resulting training dataset included a total of 2,332,722 MS^2^-biotext pairs used for training the Spec2Biotext model. For each training iteration, a single biotext was randomly sampled from the set of available biotexts for each metabolite to form the MS^2^-biotext training pair. This random sampling strategy ensures that the model is exposed to diverse functional perspectives across epochs, effectively leveraging the multi-aspect biotext annotations without modifying the standard pairwise contrastive loss formulation.

### Spec2Biotext model architecture design

To enable semantic matching between MS^2^ spectra and biotext, we developed a dual-tower deep learning architecture, Spec2Biotext. The framework consists of two modality-specific encoders: a spectral encoder for spectral input (MS^2^ spectra) and a biotext encoder for biotext snippets (biotext).

#### Spectral encoder

This encoder builds on the pretrained MSBERT backbone, which features a 6-layer Transformer encoder^74^. For each MS^2^ spectrum, we converted it to a pair of tokenized *m/z* sequences and normalized intensity values. To achieve this, we constructed a vocabulary covering the *m/z* range of 0-1000 with a resolution of 0.01 Da, supplemented with special tokens ([PAD], [MASK]). Each *m/z* value, including the precursor ion mass, was discretized to two decimal places and mapped to a token index. Intensities were normalized by dividing by the maximum peak intensity and prepended with a fixed value to preserve precursor information. Both *m/z* tokens and intensity sequences were truncated or zero-padded to a fixed maximum length (default: 100) to ensure uniform input dimensions. Only positive-ion spectra with a resolvable compound identifier and a non-missing precursor mass ≤ 1000 Da (matching the 0.00-999.99 *m/z* vocabulary) were retained; spectra missing these fields or with non-positive polarity were excluded. Peaks with *m/z* > 1,000 within otherwise valid spectra were ignored during tokenization. The model input consisted of tokenized *m/z* arrays and normalized intensity profiles. The encoder then mapped each MS^2^ spectrum to a 768-dimensional spectral embedding vector.

#### Biotext encoder

The biotext encoder employs the pretrained SPECTER model (allenai-specter; 12-layer Transformer, hidden size 768) to transform each biotext into a semantic vector representation^75^. Input texts (biotexts) were tokenized using the standard WordPiece tokenizer and truncated or zero-padded to a maximum length of 512 tokens. The [CLS] token embedding from the final encoder layer was extracted as the text-level representation. The [CLS] token is a special token placed at the beginning of the input sequence in BERT-based models, and its final hidden-state representation is commonly used as a fixed-length summary of the entire input text. This vector was subsequently passed through a linear projection layer to match the 768-dimensional latent space used by the spectral encoder. Finally, the projected embeddings were L2-normalized (scaled to unit length) to facilitate cosine-similarity-based contrastive training. The resulting fixed-length numerical vector is referred to as the biotext embedding.

#### Training

Both spectral embedding and biotext embeddings are L2-normalized (scaled to unit length by dividing by their Euclidean norm) and projected into a shared latent space (768 dimensions). To construct the validation set, for each metabolite in HMDB with multiple MS^2^ spectra, we held out one randomly selected MS^2^ spectrum as the internal validation set, while the remaining spectra were used for training. This internal split was used to monitor optimization and cross-modal retrieval during training; because the same metabolite may be represented in both training and validation through different spectra, metabolite-level generalization was assessed separately using the external and temporal holdout benchmarks described below.

### Contrastive and auxiliary objectives

Our model is trained with a cross-modal contrastive learning objective that aligns spectral embedding and biotext embedding in a shared semantic space^42^. The primary training goal is to maximize the similarity between matched MS^2^-biotext pairs and minimize similarity between mismatched pairs. We adopt an InfoNCE-based contrastive loss as the main objective^76^ and incorporate an auxiliary task: masked language modeling (MLM) on the biotext encoder^73^. The final loss is a weighted combination of these two objectives.

#### Contrastive loss

We encode the MS^2^ spectrum and its corresponding biotext using modality-specific encoders. The resulting embeddings are L2-normalized to produce **ṽ***_i_* (for MS^2^ spectrum) and **ũ***_i_* (for biotext), which lie in a shared semantic space.

We use a symmetric InfoNCE loss that encourages the similarity between matched pairs to be higher than that of unmatched ones. The contrastive objective is defined as:

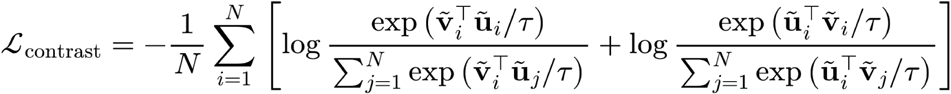

where *τ* is a temperature hyperparameter. In our study, the temperature parameter *τ* was set to 0.2. The first term maximizes the similarity between each spectrum and its corresponding text, while the second term does the same in the reverse direction.

#### Masked language modeling loss

To enhance language understanding and improve the quality of text representations, we apply a Masked Language Modeling (MLM) objective to the biotext encoder^73^. For a given input sequence x=(*x*_1_,…,*x_t_*) of *T* tokens, a subset of positions *M* ⊆ {1,….,*T*} is selected for prediction following the standard BERT-style masking strategy: 80% replaced with [MASK], 10% with random tokens, and 10% left unchanged. The model is then trained to recover the original tokens *y_t_* at masked positions. The MLM loss is defined as:

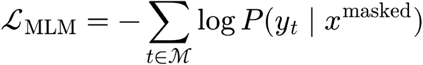

#### Total loss

The total training loss is a weighted sum of the contrastive loss and MLM loss:

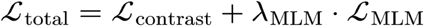

where λ_MLM_ is scalar hyperparameters that balance the auxiliary tasks. In our experiments, we set λ_MLM_=0.01. These values were kept fixed throughout training.

### Validation datasets

To evaluate the performance of the Spec2Biotext model and Spec2Function framework, we curated two independent validation MS^2^ spectral datasets, both acquired in positive ionization mode but using distinct chromatographic separation techniques: reversed-phase liquid chromatography (RPLC) and hydrophilic interaction liquid chromatography (HILIC). These datasets were generated on a Thermo Q Exactive Hybrid Quadrupole-Orbitrap Plus mass spectrometer (Thermo Fisher Scientific, San Jose, CA, USA), operated in full MS-scan mode. MS^2^ spectral data were acquired across an *m/z* range of 50 to 1000, with a scan rate of ∼4 Hz and a resolution of 30,000 at *m/z* 400. MS^2^ spectra were collected using data-dependent acquisition (DDA) of the top 10 most intense precursor ions, with normalized collision energies (NCE) set to 25 and 50 to capture fragmentation diversity^77^.

For the RPLC-based validation dataset, a total of 252 unique metabolites were detected and matched to 738 MS^2^ spectra. The HILIC-based dataset comprised 342 unique metabolites and 1,176 MS^2^ spectra. All metabolites included in both datasets were confirmed to have valid HMDB identifiers, ensuring compatibility with the Spec2Biotext and Spec2Function frameworks. These two chromatographic modes provide complementary chemical coverage and serve to benchmark the generalizability of our models across acquisition conditions and compound classes.

After merging the RPLC and HILIC datasets and removing duplicates, a total of 404 unique metabolites with valid HMDB identifiers were retained for functional inference evaluation in the Spec2Function framework.

### Benchmark methods for single-metabolite functional inference

To evaluate the functional annotation quality of Spec2Function at the single-metabolite level, we compared it against four alternative strategies that represent distinct paradigms for deriving biological descriptions from MS^2^ spectra. Their native inputs, outputs, methodological categories, and evaluation roles are summarized in Supplementary Table 1; because the native outputs differ, all methods were converted to a common functional-description output before evaluation against the same HMDB reference descriptions.

#### DeepSeek-V3

The raw MS^2^ spectrum (precursor *m/z*, fragment *m/z* values, and normalized intensities) was directly provided to a large language model (DeepSeek-V3) without any retrieval or database lookup. The model was prompted to generate a free-text functional description solely from the MS^2^ spectra, serving as a baseline to quantify how much biological knowledge an LLM can infer from mass spectral data alone.

#### Random

For each query spectrum, k metabolites (where k matched the number of candidates used by Spec2Function) were uniformly sampled at random from FemDB, irrespective of any spectral or structural similarity. The sampled metabolites and their biotexts were then fed into the same LLM prompt used by Spec2Function to generate a functional description. This baseline isolates the contribution of the retrieval component in the RAG framework.

#### GNPS

MS^2^ spectra were first matched against the GNPS spectral library using molecular networking cluster results. Matched spectra were mapped to HMDB identifiers via InChIKey. For successfully mapped metabolites, the corresponding HMDB functional description was used as the predicted output. For spectra that could not be mapped to a valid HMDB entry, a randomly selected HMDB functional description was used as a fallback.

#### CSU-MS2

The pretrained CSU-MS2 model^33^ was used to perform MS^2^-to-SMILES cross-modal retrieval following the official protocol. Candidate metabolites were filtered from HMDB by precursor neutral mass within a 10 ppm tolerance window. Spectral embeddings were compared against SMILES embeddings of all candidates, and the top-1 match was selected. The matched SMILES was mapped back to its HMDB accession, and the corresponding functional description was used as the predicted description.

#### Evaluation metric

To quantify the agreement between predicted and ground-truth functional descriptions, we adopted a text-based cosine similarity metric. Both the ground-truth functional description and the predicted text from each method were encoded using the pretrained SPECTER model (allenai-specter)^75^, an independently trained scientific language model not used in Spec2Biotext training. The [CLS] token embedding from the final Transformer layer was extracted and L2-normalized for each text. Cosine similarity was then computed as the dot product between the normalized embedding pairs. Sample-level similarities were aggregated across the validation set and compared using paired Wilcoxon signed-rank tests.

### Temporal HMDB holdout benchmark

To evaluate temporal generalization to metabolites absent from both model training and the retrieval database, we constructed a frozen historical benchmark using HMDB v4 as the knowledge boundary and metabolites newly introduced in HMDB v5 as the test set. The frozen HMDB v4 resource contained 114,222 metabolites and 519,769 BioText records, and the corresponding Spec2Biotext training set comprised 4,617 metabolites and 126,874 positive-ion MS^2^ spectra. FemDB was constructed exclusively from HMDB v4 BioTexts.

HMDB v5 metabolites were excluded if an identical metabolite or the same InChIKey connectivity was present in v4 and were further required to have available MS/MS spectra and sufficient v5 reference text. After preprocessing and harmonization across methods, the final benchmark comprised 673 v5-specific metabolites represented by 7,153 positive-ion MS^2^ spectra, with no HMDB accession overlap between the v4 training set and v5 test set.

The same five methods and evaluation procedures used in the single-metabolite benchmark were applied, with all training, retrieval, candidate-structure, and functional-text resources restricted to the frozen HMDB v4 knowledge base. For CSU-MS2 and GNPS queries without a usable frozen-v4 BioText mapping, deterministic random v4 BioText substitution was used to maintain an identical paired evaluation set across methods, and native coverage was recorded separately. Functional performance was evaluated against HMDB v5 reference BioTexts using SPECTER cosine similarity. For statistical comparisons, scores from multiple spectra of the same metabolite were first aggregated at the metabolite level, followed by paired Wilcoxon signed-rank tests.

### Robustness evaluation across instrument platforms and biological matrices

#### Cross-instrument holdout evaluation

Instrument annotations for MassSpecGym spectra were restored from the original records, and spectra without defined instrument families were excluded. Two symmetric leave-instrument-out models were trained: QTOF spectra were excluded and the model was trained on 91,611 Orbitrap spectra, or Orbitrap spectra were excluded and the model was trained on 35,385 QTOF spectra. Evaluation was restricted to 2,728 HMDB compounds measured on both platforms in positive mode with the same adduct, with one matched QTOF–Orbitrap pair selected per compound. The corresponding same-compound spectrum from the training platform was permitted during training; thus, this analysis assessed cross-instrument transfer rather than generalization to unseen metabolites. Held-out spectra were analyzed using Spec2Function, direct DeepSeek-V3 prompting, or random FemDB retrieval, and generated annotations were compared with HMDB BioTexts using SPECTER cosine similarity. Paired one-sided Wilcoxon signed-rank tests were used for comparisons with the two baselines.

#### Paired biological and reference spectrum comparison

MS/MS spectra from feces, plasma, saliva, and urine were searched against the frozen Spec2BioText training spectral library using metid::match_ms2_temp() (metid v1.3.2), using a 25-ppm precursor window and the default 30-ppm fragment tolerance. Top-ranked matches passing the default search cutoff (SS > 0.50) were initially retained, and the primary analysis was restricted to high-confidence pairs with SS ≥ 0.70, yielding 55 spectrum pairs representing 42 unique HMDB compounds. Biological and matched reference spectra were independently processed through the same Spec2Function pipeline using the top five unique FemDB entries and DeepSeek-V3 (temperature = 0.2), while the matched compound identity and HMDB BioText were withheld until evaluation. Annotation quality was measured against the corresponding HMDB BioText using SPECTER cosine similarity. Scores were averaged by HMDB compound when necessary and compared using a two-sided Wilcoxon signed-rank test; 95% confidence intervals were estimated by compound-clustered bootstrap resampling.

#### Controlled spectral degradation analysis

Spectral-quality robustness was evaluated using 404 molecule-unique spectra from the external RPLC/HILIC benchmark. Starting from the original spectra, four types of degradation were independently introduced: fragment-peak deletion (10%, 25%, 50%, and 75%), fragment-intensity perturbation (σ = 0.10, 0.25, and 0.50), addition of background peaks (10%, 25%, 50%, and 100% of the original fragment count), and chimeric mixing with an unrelated spectrum (10%, 25%, and 50% donor signal). A precursor-only condition, generated by removing all fragment peaks, was included as a negative control. Perturbed spectra were processed through the same frozen Spec2Function workflow used for the external benchmark. Functional annotation quality was assessed by SPECTER cosine similarity to the corresponding HMDB BioText. The predefined spectrum-to-FemDB cosine threshold of 0.65 was retained as the evidence-support gate, and robustness was summarized by both annotation similarity among retained predictions and the fraction of spectra remaining above this threshold.

### Leakage-controlled and frequency-normalized PubMed literature validation

To assess literature support for functional modules identified by Spec2Function and pathways identified by comparator workflows while minimizing knowledge-source leakage and publication-frequency bias, we implemented a leakage-controlled and frequency-normalized PubMed validation procedure using NCBI E-utilities (esearch.fcgi).

For each Spec2Function-derived functional module, all PubMed records retrieved and used as evidence during module-level functional inference were recorded by PubMed identifier (PMID). These publications were treated as the module-specific exclusion set and were removed from all subsequent literature-validation queries for that module, including both searches for the functional term alone and searches for the functional term in combination with the biological phenotype. Thus, publications used to generate the functional interpretation could not also contribute to its subsequent literature validation. For pathways obtained from comparator workflows that did not use PubMed literature during pathway generation, the corresponding exclusion set was empty.

For each pathway or functional module term (A), PubMed was queried separately for the term alone and for its co-occurrence with the phenotype of interest. For the liver cirrhosis analysis, the phenotype query was “liver cirrhosis”, whereas aging analyses used “aging” together with the corresponding tissue term where applicable. Composite module names were mapped to their corresponding parent biological process or pathway term when necessary to ensure consistent querying.

Literature support was normalized by the overall publication frequency of each functional term and calculated as

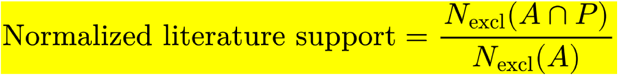

where *P* denotes the phenotype context, *N*_excel_(*A*) is the number of PubMed publications associated with term *A* after removal of the module-specific exclusion set, and *N*_excel_(*A*∩*P*) is the number of remaining publications in which term *A* and phenotype *P* co-occur. Terms with no remaining background publications were not assigned a normalized score.

The normalized literature-support score, rather than the absolute PubMed publication count, was used for comparisons between Spec2Function-derived functional modules and pathways identified by conventional workflows. Raw publication counts were retained only as descriptive information.

### Functional embedding metabolite database (FemDB)

To support functional similarity search within the Spec2Function framework, we constructed the Functional Embedding Metabolite Database (FemDB). Starting from the full set of 248,138 metabolites in HMDB (version 5.0 retrieved on March 18, 2025) with valid identifiers and curated biological annotations, we applied the standardized template-based biotext generation pipeline described above to produce biotexts for each compound, covering molecular identity, taxonomic classification, biological localization, biospecimen distribution, pathway and protein interactions, disease associations, and biological processes.

Each biotext was then passed through the biotext encoder of the trained Spec2Biotext model to produce a fixed-length vector representation, referred to as a functional embedding. These embeddings capture the latent biological semantics of each metabolite and are indexed together to form FemDB. In total, FemDB comprises 2,515,806 functional embeddings derived from 248,138 HMDB metabolites, with multiple embeddings per metabolite reflecting distinct functional aspects. FemDB serves as the reference search space for embedding-based retrieval throughout Spec2Function, enabling rapid and scalable identification of biologically similar metabolites to an input MS^2^ spectrum without requiring structural identity or direct spectral matches.

### Spec2Function framework design

The Spec2Function framework supports functional annotation of untargeted MS^2^ spectra through two computational modes: (i) single-MS^2^ spectrum functional inference and (ii) metabolite set functional annotation.

#### Mode 1

In Mode 1, the input is a single MS^2^ spectrum. This MS^2^ spectrum is first preprocessed and converted into spectral embedding using the spectral encoder trained within the Spec2Biotext model. The resulting spectral embedding is queried against the FemDB. Cosine similarity is used to retrieve the top functionally relevant matches. A fixed evidence threshold of 0.65 is applied before downstream functional inference. If no FemDB entry reaches this threshold, the query returns “insufficient evidence” and no functional description is generated. Otherwise, metabolites above the threshold are ranked by similarity and the top 5 are retained as functionally related evidence. These retrieved metabolites are then passed into the functional inference module for downstream functional annotation of the input MS^2^ spectrum.

#### Mode 2

In this mode, Spec2Function performs set-level functional annotation. The input is a collection of MS^2^ spectra corresponding to differentially expressed metabolic peaks, optionally accompanied by pre-existing structural annotations. If structural annotations are already available (*e.g.*, obtained via conventional MS^2^ spectral library matching using tools such as the metid package^77^ from the TidyMass project^7,8^), these annotated metabolites are directly incorporated into the analysis. In our liver cirrhosis case study, structural annotation was performed using the metid package prior to Spec2Function analysis. Both structurally annotated metabolites and functionally similar metabolites retrieved from FemDB are aggregated into a unified metabolite set, which is then submitted to the functional inference module to synthesize the biological functions of the input metabolite set.

### Function inference module

For each metabolite involved in the analysis, whether obtained through MS^2^ spectra-based structural annotation or embedding-based functional similarity retrieval, the module first retrieves relevant literature from the PubMed database. Literature retrieval is performed using metabolite names and their known synonyms as search queries, and only the title and abstract of each matched publication are retained to ensure concise and interpretable input. In parallel, the curated biotext generated from the Human Metabolome Database (HMDB) is retrieved for each metabolite from the functional embedding database (FemDB), providing structured biological context including pathway associations, disease relevance, and tissue localization.

#### Mode 1 (single-spectrum inference)

For a query MS^2^ spectrum, the encoded embedding is used to retrieve the top-k most functionally similar HMDB entries from FemDB by cosine similarity. The top 5 retrieved metabolite names are then used to query PubMed for supporting literature. The retrieved HMDB biotexts and PubMed abstracts are assembled into a standardized expert-level prompt and passed to the LLM (gpt-4o), which is instructed to act as a domain expert in biochemistry and molecular biology and to produce a biological function summary for the query peak in a scientific, concise, and non-redundant manner. The output covers primary biological classification, pathway involvement, disease relevance, and an overall confidence assessment.

#### Mode 2 (set-level inference)

For a metabolite set, function inference operates at the cluster level following unsupervised spectral clustering. For each cluster, functional evidence is retrieved from FemDB using one of two strategies: centroid-based retrieval, in which the cluster centroid embedding is queried against FemDB to identify the top-k most similar metabolites; or voting-based retrieval, in which each spectrum in the cluster independently retrieves its top-k FemDB matches, and results are aggregated by vote frequency and maximum similarity across all cluster members. The resulting ranked metabolite list, combined with HMDB biotexts and cluster-specific PubMed literature retrieved via an LLM-generated search query, is passed to the LLM in two sequential calls: the first generates a KEGG-style functional theme name for the cluster, and the second produces a process-focused biological report. After all clusters are processed, a second-tier LLM call synthesizes a global biological narrative integrating the functional themes and supporting evidence across all clusters.

A key feature shared across both modes is a similarity-weighted aggregation strategy. For metabolites retrieved via functional embedding similarity, the cosine similarity score to the query MS^2^ spectrum serves as the confidence weight. For metabolites structurally annotated via MS^2^ spectral matching, the MS^2^ match score is used instead. These weights are explicitly incorporated into the LLM prompt: retrieved metabolites are ranked by score, confidence tiers are labeled (High: ≥ 0.85; Medium: 0.65-0.85; Abstain: < 0.65), and higher-confidence entries receive proportionally more detailed biotext input. The model is explicitly instructed to prioritize higher-confidence evidence in its biological reasoning. This design reduces the risk of overinterpreting weakly relevant annotations and enhances the biological specificity of the generated summaries.

### Evaluation of functional similarity captured by biotext embeddings

To assess whether Spec2Biotext embeddings capture biochemical relatedness, cosine similarity was computed between metabolite pairs with shared or distinct pathway and chemical class annotations derived from HMDB. Biochemically related pairs were defined as metabolites sharing both pathway membership and chemical subclass. Biochemically unrelated pairs were defined as metabolites sharing neither pathway membership nor chemical superclass. For each category, 10,000 pairs were randomly sampled. Biotext embeddings were obtained from FemDB, where each metabolite is represented by a 768-dimensional vector generated by Spec2Biotext. Only metabolite pairs with embeddings available for both metabolites were retained. Cosine similarity was calculated for each pair, and similarity distributions between positive and negative pairs were compared using a two-sided Wilcoxon rank-sum test.

### Evaluation of pathway-level functional inference

To evaluate Spec2Function’s ability to infer biological functions at the pathway level, we selected pathways from the PathBank database^78^ using two inclusion criteria: all metabolites in the pathway had corresponding HMDB MS/MS spectra, and each pathway contained at least two metabolites. This yielded 27 pathways for evaluation. Ground-truth functional descriptions were taken directly from PathBank pathway entries. For each pathway, the MS^2^ spectra of all constituent metabolites were encoded using the spectral encoder and aggregated into a single pathway-level representation by computing the mean embedding vector, followed by L2 normalization (centroid aggregation). This pathway centroid was then used to query FemDB and retrieve the top functionally similar metabolites for downstream LLM-based functional annotation using the Mode 2 inference module. A random control was generated by permuting MS^2^ spectra across pathways: for each pathway, spectra were drawn at random from the full validation set (matching the original pathway size), and the same centroid aggregation and annotation procedure was applied. Functional annotation quality was evaluated using cosine similarity between embeddings of the predicted description and the PathBank ground-truth description, computed using the SPECTER encoder. Differences between Spec2Function and the random baseline were assessed using the Wilcoxon signed-rank test.

### Biofluid-specific functional profiling from representative LC-MS/MS peaks

Publicly available untargeted LC-MS/MS metabolomics datasets from four human biofluids (plasma, urine, feces, and saliva) were obtained from the MassIVE repository. Raw data were processed using the TidyMass workflow in positive-ion mode^7,8^. Metabolic peaks were detected and aligned using a mass tolerance of 10 ppm and a peak width of 10–60 s. Peaks with missing values in more than 50% of samples were removed; remaining missing values were imputed using k-nearest neighbors followed by median normalization. MS^2^ spectra were matched to metabolic peaks using a precursor mass tolerance of 10 ppm and a retention time tolerance of 15 s. Peaks with confident MS^2^-based annotations from reference spectral libraries were excluded. Spectra containing fewer than five fragment ions were removed. The remaining unannotated MS^2^-associated peaks were ranked by mean intensity within each dataset, and the top 100 were retained as representative biofluid features.

For each of the 100 representative MS^2^ spectra per biofluid, a per-spectrum functional annotation was generated using Spec2Function Mode 1. To aggregate annotations across spectra, each LLM-generated description was mapped to a set of predefined functional categories through keyword matching. A curated dictionary of 57 keywords was used to assign each annotation to one or more of 15 biological functional categories (*e.g.*, microbial metabolism, xenobiotic metabolism, oxidative stress, lipid metabolism, inflammation; full list in Supplementary Data 7). For each biofluid, the number of spectra mapped to each functional category was tallied, and the counts were normalized by the total number of annotated spectra in that biofluid to obtain a proportion value (0-1). Functional categories with a maximum proportion below 0.01 across all biofluids were excluded.

### Comparative analysis of Spec2Function and alternative workflows in liver cirrhosis

The liver cirrhosis plasma metabolomics dataset was obtained from the MetaboLights repository (accession: MTBLS5665)^51^. Raw LC-MS/MS data were processed using the TidyMass ecosystem^7,8^. For the identification-dependent workflow, metabolite annotation was first performed using reference MS^2^ spectral libraries, and only level 2 annotations were retained. When multiple annotations were assigned to the same peak, the candidate with the highest spectral similarity score was selected. Differential analysis was then conducted at the metabolite level using identified metabolites, with statistical significance assessed by two-sample t-tests followed by Benjamini-Hochberg correction. Metabolites with FDR < 0.05 were considered significantly altered and were used for KEGG pathway enrichment analysis. For the Spec2Function workflow, differential analysis was performed independently at the LC-MS/MS peak level. Peaks with FDR < 0.05 were considered significant, and their corresponding MS^2^ spectra were extracted for downstream analysis. All differential MS^2^ spectra without annotation, including those derived from structurally unannotated peaks, were used as input for Spec2Function Mode 2 to infer biological functions directly from spectral data.

For the CSI:FingerID benchmark, the same 442 differential MS^2^ features were analyzed using SIRIUS/CSI:FingerID v6.3.4 with the Orbitrap profile and BIO structure database^55^. Top-ranked structure candidates were ordered by ConfidenceScoreExact, and the highest-confidence 10% of successfully annotated features were retained. Candidates were collapsed by two-dimensional InChIKey connectivity and mapped to human KEGG compound identifiers. Pathway over-representation analysis was performed using a one-sided hypergeometric test followed by Benjamini–Hochberg correction.

For the MetDNA benchmark, the MTBLS5665 LC-MS data were reconstructed using XCMS and analyzed using MetDNA v1.1.2^56^. The 442 differential features were matched to reconstructed LC-MS features using precursor m/z and retention time, followed by spectral-library-based seed annotation and metabolic reaction-network propagation. The top-ranked KEGG annotation for each matched feature was retained, and unique compounds mapping to the human KEGG pathway background were analyzed using the same KEGG compound–pathway database, hypergeometric test, and Benjamini–Hochberg procedure used for CSI:FingerID.

For the MS2MP benchmark, the same 442 differential MS^2^ features were evaluated using the official MS2MP workflow^19^. SIRIUS v5.6.3 was first used to generate the fragmentation trees required by MS2MP. Of the 442 differential spectra, 389 generated MS2MP-compatible fragmentation trees and output records, of which 278 received at least one pathway label. Dataset-level pathway enrichment was performed using the MS2MP-compatible full-feature background and a one-sided hypergeometric test, followed by Benjamini–Hochberg correction across model-emitted pathway labels. Because MS2MP directly outputs pathway labels rather than metabolite identities, its native feature-level pathway framework was retained rather than remapping predictions to the compound-level human KEGG background used for CSI:FingerID and MetDNA.

### Multi-organ rhesus macaque aging metabolomics analysis

Untargeted LC-MS/MS metabolomics data were obtained from the published multi-omics aging atlas of female rhesus macaques, comprising 30 tissues from 17 animals aged 3-27 years^57^. For the identification-dependent workflow, the data were processed with TidyMass for peak extraction and MS^2^-based metabolite annotation, and differential analysis was performed on the annotated metabolites. Within each tissue, log2-transformed peak areas were modeled as a function of age (molecular abundance ∼ age) and analyzed with empirical Bayes moderation in limma, following the original study. Metabolites with |β| > 0.008 and *P* < 0.05 were considered age-associated. Annotated metabolites with valid KEGG compound identifiers were used for KEGG pathway analysis. The full results are provided in Supplementary Data 8.

For the Spec2Function workflow, the same analysis was applied directly to all peaks with MS^2^ spectra. Within each tissue, log2-transformed peak intensities were modeled against age using the same limma-based framework. The age-associated spectra without annotation were then analyzed with Spec2Function Mode 2 using centroid-based FemDB retrieval (top-k = 5; minimum cosine similarity = 0.65), followed by functional module generation with the two-tier LLM inference strategy described above.

### Cross-species validation of age-associated functional modules

To assess whether Spec2Function-inferred macaque modules were reproducible across species, we validated them against two independent publicly available aging metabolomics datasets. For mouse validation, metabolite abundance data from a 10-organ aging metabolomics atlas (Mus musculus; MetaboLights accession: MTBLS10642) were analyzed across four age groups (4, 8, 12, and 20 months) using one-way ANOVA with Benjamini-Hochberg correction to identify age-associated metabolites. Spearman’s rank correlation was used to characterize monotonic age-related trends. KEGG over-representation analysis was then performed per organ on the significant metabolites. For human validation, a plasma metabolomics dataset comparing young and elderly subjects (MetaboLights accession: MTBLS266) was analyzed using two-sample t-tests followed by KEGG over-representation analysis.

To characterize the relationship between Spec2Function discoveries and established aging metabolic signatures, identified modules were classified into two categories: Consensus modules, detected by both Spec2Function and conventional KEGG enrichment in macaque tissues (FDR < 0.05), and Spec2Function-specific modules, identified by Spec2Function but not by conventional KEGG enrichment under the analysis settings used here. Cross-species reproducibility was assessed by examining whether the corresponding KEGG pathways were enriched in the mouse and human aging datasets.

## Code availability

All software development, data processing, and statistical analyses were performed using Python (version 3.10.18) and R (version 4.4.1) along with their respective packages. The full source code for Spec2Function is openly available at https://github.com/jaspershen-lab/spec2function. The online interactive version of Spec2Function is accessible at https://spec2function.jaspershenlab.com. A dedicated reproduction repository containing benchmarking and validation scripts, computational environment specifications, and analysis workflows used in the revised manuscript is available at https://github.com/jaspershen-lab/Spec2Function_reproduction.

## Data availability

All datasets analyzed in this study and used as demonstration data were obtained from previously published sources^25,38–41^. Additional source data supporting the findings of this study are provided in the Supplementary Data. The human biofluid LC-MS/MS datasets used in this study are available through the MassIVE repository (https://massive.ucsd.edu/ProteoSAFe/) under the following accessions: plasma (MSV000090524), urine (MSV000082261), feces (MSV000095787), and saliva (MSV000083049). The liver cirrhosis metabolomics dataset used for case analysis is available in the MetaboLights repository (https://www.ebi.ac.uk/metabolights/MTBLS5665). Raw LC-MS/MS metabolomics data from the rhesus macaque multi-organ aging atlas are available in OMIX (https://ngdc.cncb.ac.cn/omix, accession no. OMIX001779). The Functional Embedding Metabolite Database (FemDB) used by Spec2Function is freely available through Hugging Face at https://huggingface.co/cgxjdzz/ms2function-assets. The required FemDB assets are automatically downloaded from this repository when running Spec2Function and do not require separate manual installation.

## Acknowledgments

This work was supported by start-up funding (Nanyang Assistant Professorship) awarded to Xiaotao Shen from the Lee Kong Chian School of Medicine and the School of Chemistry, Chemical Engineering, and Biotechnology at Nanyang Technological University, Singapore. This work was also supported by a Tier 1 grant from the Ministry of Education, Singapore, to Xiaotao Shen (Grant number: #025402-00001). Chuchu Wang was also supported by start-up funding (Nanyang Assistant Professorship) from LKCMedicine at NTU Singapore. The authors acknowledge the use of ChatGPT (OpenAI) for assistance with language polishing and grammar correction during the preparation of this manuscript.

## Author contributions

X.S. and F.Z. jointly conceived the project. X.S. supervised the overall study. F.Z. and X.S. designed the methodology. F.Z. and Y.L. prepared the MS^2^ spectra library. F.Z. developed the software. Case study analyses were performed by F.Z., Y.W., and C.W.. Y.W., X.S., and F.Z. designed and created the atlas website. X.S., F.Z., Y.W., and C.W. prepared all figures. The manuscript was written by X.S., F.Z., and Y.W., with all authors reviewing and contributing to the final version.

